# Creatine utilization as a sole nitrogen source in *Pseudomonas putida* KT2440 is transcriptionally regulated by CahR

**DOI:** 10.1101/2021.11.02.466972

**Authors:** Lauren A. Hinkel, Graham G. Willsey, Sean M. Lenahan, Korin Eckstrom, Kristin C. Schutz, Matthew J. Wargo

## Abstract

Glutamine amidotransferase-1 domain-containing AraC-family transcriptional regulators (GATRs) are present in the genomes of many bacteria, including all *Pseudomonas* species. The involvement of several characterized GATRs in amine-containing compound metabolism has been determined, but the full scope of GATR ligands and regulatory networks are still unknown. Here, we characterize *Pseudomonas putida’*s detection of the animal-derived amine compound, creatine, a compound particularly enriched in muscle and ciliated cells by a creatine-specific GATR, PP_3665, here named CahR (<u>C</u>reatine <u>a</u>mido<u>h</u>ydrolase <u>R</u>egulator). *cahR* is necessary for transcription of the gene encoding creatinase (*PP_3667/creA*) in the presence of creatine and is critical for *P. putida*’s ability to utilize creatine as a sole source of nitrogen. The CahR/creatine regulon is small and electrophoretic mobility shift demonstrates strong and specific CahR binding only at the *creA* promoter, supporting the conclusion that much of the regulon is dependent on downstream metabolites. Phylogenetic analysis of creA orthologs associated with cahR orthologs highlights a strain distribution and organization supporting likely horizontal gene transfer, particularly evident within the genus *Acinetobacter*. This study identifies and characterizes the GATR that transcriptionally controls *P. putida* metabolism of creatine, broadening the scope of known GATR ligands and suggesting GATR diversification during evolution of metabolism for aliphatic nitrogen compounds.

## INTRODUCTION

Like many primarily soil-dwelling microbes, the Gram-negative bacterium *Pseudomonas putida* has evolved a vast and diverse array of transport and metabolic machinery to fuel its organoheterotrophic lifestyle ^(1)^. Within the rhizosphere, fast and efficient adjustment to the surrounding environment ensures successful acquisition of essential nutrients and activation of stress responses that are crucial to *P. putida*’s survival ^(2)^. Effective response to the extracellular environment is partly afforded by the large number of transcription regulatory proteins encoded by the *P. putida* genome. About 2% of the predicted genes in the 6.15 Mbp *P. putida* KT2440 genome contain a conserved AraC-type DNA-binding Helix-Turn-Helix motif, characteristic of many catabolism-related transcription regulators ^(3)^. These AraC-family transcription regulators control a large number of metabolic processes within *P. putida*, although for many, the cognate inducing ligands and target regulons have not been identified.

Creatine is an amine compound found primarily in animal tissues where it serves to buffer charging of high-energy carriers during rapid ADP to ATP conversion. It is particularly abundant in tissues that require large pools of ATP and ADP for periods of intense energy expenditure, such as fast twitch skeletal muscle and ciliated cells ^(4)^. Metabolism of dietary creatine in animals occurs via partial metabolism and modification by gut-residing bacteria creating 1-methylhydantoin, a metabolite that is not easily metabolized by the body and can lead to tissue inflammation. To prevent creation and accumulation of 1-methylhydantoin, the major avenue of creatine homeostasis in animals is the excretion of creatine and creatinine in the urine, where it can be used by soil microbes. *Pseudomonas* species have been observed to metabolize creatine and creatinine, as well as the microbial breakdown product, 1-methylhydantoin ^(4), (5), (6), (7), (8), (9), (10), (11), (12)^. The products of creatine and creatinine metabolism by *Pseudomonas* vary depending on the specific degradation pathway used, which is species and sometimes strain dependent. For creatinine, the process often begins with breakdown of creatinine to creatine by a creatininase (creatinine amidohydrolase), though there are alternate pathways of metabolism. Creatine is then further metabolized to sarcosine and urea by creatinase (creatine amidinohydrolase), and sarcosine is then converted to glycine and formaldehyde by the tetrameric sarcosine oxidase (SoxBDAG) (**Fig. 1A**), or to methylamine by sarcosine reductase and/or glycine reductase ^(4), (5), (13), (14)^.

**Figure 1.**
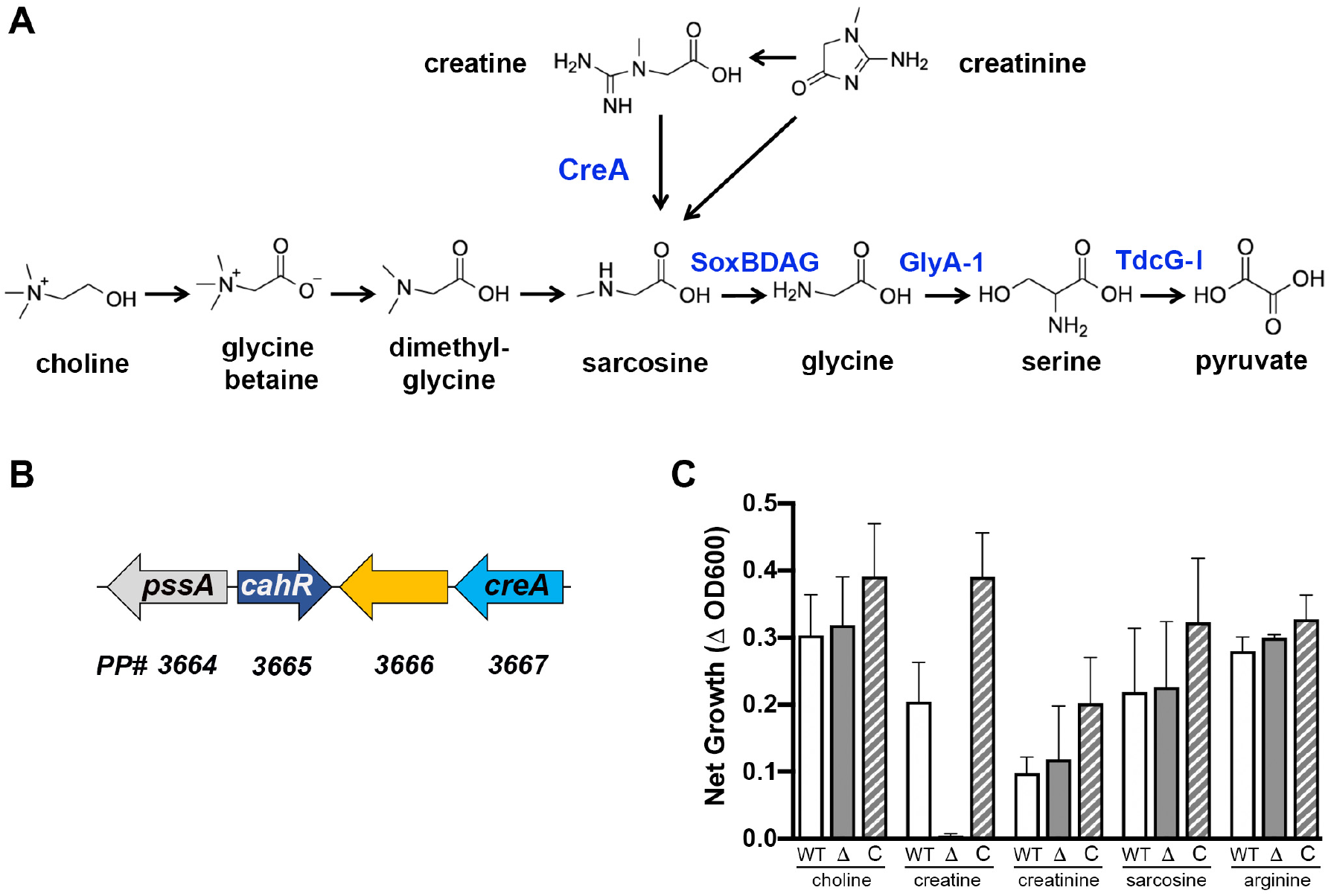
The creatine metabolic pathway and the role of *PP_3665/cahR* in creatine metabolism. **(A)** The creatine metabolic pathway in *P. putida* intersects with the choline metabolic pathway at sarcosine. The enzymes primarily discussed in this study are noted in blue. **(B)** Genomic organization around the creatinase gene (*PP_3667/creA*) in *P. putida* KT2440. Gene colors match those used in Figure 6. **(C)** Net growth of *P. putida* wild type (WT), Δ*PP_3665/cahR* (Δ), and the complemented strain (C), compared to nitrogen free minimal media controls. The compounds present as the sole nitrogen source are listed below each group of bars. Error bars denote standard deviation from between three and six experiments.

Production of creatinase is inducible in *P. putida* when the bacterium is supplied with creatine as the sole carbon or nitrogen source, but the induction mechanism is currently unknown ^(8)^. *P. putida PP_3665* encodes an AraC-type transcription regulator convergently transcribed with an operon containing the creatinase gene, *creA* (**Fig. 1B**). Among characterized *Pseudomonas* genomes, only a few, including particular strains of *P. putida* and *P. resinovorans*, maintain likely orthologs of both the creatinase gene and the associated AraC-family transcription regulator in a similar syntenic arrangement ^(3)^. Both of these species of *Pseudomonas* are present primarily in soils and are active participants in the rhizobiome, i.e. bacteria within the rhizosphere. It is within the rhizosphere that *P. putida* most likely encounters creatine, as creatine and its anhydrous form, creatinine, are present in soils, deposited via animal urine or feces as well as through animal tissue degradation ^(15), (16), (17), (18)^.

Here, we describe identification of a regulator, PP_3665, required for transcriptional induction of creatinase in *P. putida* in response to creatine and utilization of creatine as a sole nitrogen source, leading us to name PP_3665 as the <u>C</u>reatine <u>a</u>mido<u>h</u>ydrolase <u>R</u>egulator, CahR.

## MATERIALS AND METHODS

### Bacterial Strains and Growth Conditions

*Pseudomonas putida* strain KT2440 and mutants made from this parent strain were grown at 30 °C shaking (170 rpm) in morpholinopropanesulfonic acid (MOPS) media ^(19)^ with modification we previously reported ^(20)^, supplemented with 25 mM pyruvate, 5 mM arginine, and 20 μg/ml gentamicin when needed for plasmid maintenance. *Escherichia coli* strains DH5α and *S17λpir*, used for cloning and conjugation with *P. putida*, respectively, and *E. coli* strain T7 used for recombinant protein expression, were maintained in lysogeny broth (LB), Lennox formulation, supplemented with 7 μg/ml gentamicin or 150 μg/ml carbenicillin as appropriate. *E. coli* strains were grown at 37°C with shaking (170 rpm).

### Construction of the *P. putida ΔPP_3665* deletion strain and complementation construct

A 980 base pair (bp) fragment upstream of the coding region of *cahR* was amplified with primers PP3665KO_F1_HindIII (5’-AAGCTTCGGGATCGTTCCAGATGCGT-3’) and PP3665KO_R1 (5’-GATCCAGGTGCTGCCCGATGCCA-3’). A 1 kb fragment downstream of the coding region of *cahR* was amplified with primers PP3665KO_F2 (5’- GTTGGGTCAGGTATTGGCATTG-3’) and PP3665KO_R2_EcoRI (5’- GAATTCCCGAGGCGGAAAACCCCTGT-3’). The two fragments flanking the *cahR* coding region were spliced together via splice-overlap extension (SOE) PCR using primers PP3665KO_F1_HindIII and PP3665KO_R2_EcoRI, digested, and ligated into similarly cut plasmid pMQ30^(21)^ containing a gentamicin resistance cassette for initial selection and the *sacB* gene for counter selection. pMQ30 containing the flanking regions of *cahR* (pMQ30:*cahR-*KO) was maintained in *E. coli* strain DH5α, and transformed into *E. coli* S17λpir for conjugation with *P. putida*. Single crossover integrants were selected by gentamicin resistance (50 μg/ml) and double crossover to deletion or revertant was carried out on LB agar with no salt and amended with 10% sucrose, as previously described^(22), (23)^, to yield an unmarked deletion of *PP_3665/cahR* in *P. putida* strain KT2440 (strain LAH111). Deletion or reversion was determined using primers PP3665 KO screen F (5’- ATTTCACCACCATCGGCCTT-3’) and PP3665 KO screen R (5’- AGCGGTAGCCTTTGAGCAAT-3’), which yields a 3.3 kb product in WT and a 2 kb product in the deletion strain.

*cahR* complementation in the *ΔcahR* strain was achieved by plasmid expressed *cahR*. Briefly, the coding region of *cahR* and divergently transcribed gene *pssA* (to ensure inclusion of the full coding region and promoter region of *cahR*) were amplified using primers PP_3665-EXP-F_EcoRI (5’-ATAGAATTCGACATCAATGCGCGGTGC-3’) and PP_3665-EXP-R-HindIII (5’-AAAAAGCTTCTGCACTGGCTTCTTCTCAC-3’). The ~2.5 kb fragment was digested with the aforementioned enzymes and ligated into the similarly cut pMQ80 plasmid. The resulting pMQ80:*pssA/cahR* complementing plasmid was then electroporated into *P. putida* KT2440 *ΔcahR* as previously described ^(22)^.

### Nitrogen source growth assays

*P. putida* KT2440 wild type and □*cahR* strains carrying the pMQ80 empty vector, and *P. putida* KT2440 Δ*cahR* complemented with pMQ80:*pssA/cahR* were grown overnight at 30 °C shaking in 1x MOPS buffer amended with 25 mM pyruvate, 5 mM arginine, and 20 μg/ml gentamicin. Cells from overnight culture were collected by centrifugation, washed with 1x MOPS media lacking nitrogen (MOPS no nitrogen), and adjusted in 1x MOPS no nitrogen to an optical density at 600 nm (OD_600_) of 0.5. These normalized cell suspensions were added to pre-warmed 1x MOPS no nitrogen media amended with 20 mM pyruvate, 20 μg/ml gentamicin, and 2 mM of one of the following nitrogen sources: choline, creatine, creatinine, sarcosine, or arginine. A control for residual growth in media lacking a nitrogen source was also included. Cells were added to each condition to a final optical density at 600 nm (OD_600_) of 0.05 in a 500 μl final volume in wells of a 48-well plate. Cultures were incubated at 30 °C with horizontal shaking at 170 rpm for 18 hours. The optical densities of cultures were measured at times 0 hours and 18 hours using a Synergy H1 plate reader (BioTek, Winooksi, VT). Growth was reported as the net growth of each strain amended with a nitrogen source after subtraction of the optical density in the 0 mM nitrogen condition, as there is always a small amount of residual growth from stored intracellular nitrogen.

### *cahR*-dependent transcriptional induction assays and *PP_3667/creA* promoter mapping

Transcriptional reporter fusions of the full *creA* promoter, −10 bp to −195 bp from the predicted *creA* transcriptional start site, and a promoter truncation including −10 bp to −169 bp from the predicted *creA* transcriptional start site, to the coding region of GFP were constructed following a protocol for creation of recombinant plasmids similar to that described by Bryksin and Matsumura ^(24)^. The full promoter region of *creA* was amplified using *PP_3667*PromF (5’- CGGGTACCGAGCTCGTTCAGGCCGGCCGC-3’) and *PP_3667*PromR (5’-TAAGATTAGCGGATCAGACTTTGTGGC-3’) primers, and the - 169 truncation fragment was amplified using primers *PP_3667*PromF (5’- CGGGTACCGAGCTCGTTCAGGCCGGCCGC-3’) and *PP_3667*Prom-169R (5’- CGGGTACCGAGCTCGTCATGGAGCTGGACC-3’). Primers for amplification of the full and −169 truncated *creA* promoters contain 5’-regions complementary to the sequence upstream of GFP’s coding region on plasmid pMQ80. The promoter fragments flanked with pMQ80 complementary ends were mixed with plasmid pMQ80, and the mixtures were denatured, annealed and amplified via PCR to allow for insertion of the promoter region upstream of pMQ80’s GFP. Insertion of the *creA* promoter fragments resulted in deletion of pMQ80’s NheI and EcoRI restriction sites, so the amplified plasmid pool was digested with EcoRI and NheI to cull pMQ80 that did not incorporate the *creA* promoters. Following separation and purification of the circular plasmid via gel electrophoresis, the *creA* promoter-containing plasmids were transformed into *E. coli* DH5α. The −56 *creA* promoter truncation was constructed via three sequential PCRs, using the reverse primer pMQ80_GFP_R (5’-TCAGGCTGAAAATCTTCTC-3’) and the following forward primers for a 5’-extension of GFP including the truncated promoter of *creA*; PP3667-56_1 (5’- CTTCTCAGGCGGCCGGCCTGAACCGAGCTCGGTACCCG-3’), PP_3667-56_2 (5’-GTCGGTTCTGTTGCAATGCTTCTCAGGCGGCCGGCC-3’), and PP_3667- 56_3_BamHI (5’-AATGGATCCGTGTCCTCGTCGGTTCTG-3’). This −56 *creA* promoter-GFP fusion fragment was digested with restriction enzymes BamHI and HindIII and ligated into similarity cut pMQ80. The −80 *creA* promoter-GFP fusion fragment was created by further extending the −80 *creA* promoter-GFP fragment using primers pMQ80_GFP_R and PP3667-80_NheI (5’-AATGCTAGCATTCCAGGTCGGATAGATACAAAAGTGTCCTCGTCGGT-3’). The −80 *creA* promoter-GFP fusion fragment was digested with NheI and HindIII restriction enzymes and ligated into similarly cut pMQ80. All plasmids were propagated in *E. coli* DH5α, purified, and electroporated into *P. putida* KT2440 for induction assays.

Each plasmid encoding the *creA* promoter-*gfp* fusion, full or truncated, was electroporated into *P. putida* KT2440 wild type or Δ*cahR* for transcriptional induction assays. To test metabolite-specific induction using the full *creA* promoter *gfp* reporter construct, *P. putida* KT2440 wild type and Δ*cahR* carrying the pMQ80:*creA-_195_gfp* plasmid were grown overnight in 1x MOPS media amended with 25 mM pyruvate, 5 mM arginine, and 20 μg/ml gentamicin. Overnight cultures were adjusted to a uniform OD_600_ and added to 1xMOPS media amended with 20 mM pyruvate, 20 μg/ml gentamicin, and +/− 2 mM of each nitrogen containing compound to a final OD_600_ of 0.5 in a 48-well plate. The compounds creatine, creatinine, sarcosine, and glycine betaine were tested for their *cahR*-dependent induction via the *creA* promoter. Plates were incubated at 30 °C with periodic shaking for 18 hours with readings of the OD_600_ and GFP fluorescence (Excitation: 485 nm/Emission: 528 nm) taken every hour.

To determine the region of the *creA* promoter essential for creatine-responsive *cahR*-dependent induction, induction assays using *creA-gfp* transcriptional reporters engineered with truncated regions of the *creA* promoter upstream of *gfp* were conducted. The assays were conducted as described above with the following adjustments; *P. putida* KT2440 wild type strains carrying the pMQ80:*creA-_195_-gfp*, pMQ80:*creA-_169_-gfp*, pMQ80:*creA-_80_-gfp*, or pMQ80:*creA-_56_-gfp* plasmids were induced in 1xMOPS media amended with 20 mM pyruvate, 20 μg/ml gentamicin, and with or without 2 mM creatine. The full (−195) and −169 *creA* promoters were tested for *cahR*-dependence by measuring the transcriptional induction of *gfp* carried on the respective reporter plasmid in the *P. putida* KT2440 Δ*cahR* background. Fold induction was reported using the equation: Fold Induction = (Fluorescence Units_2mM N-source_/Fluorescence Units_0mM nitrogen_).

### MBP-CahR fusion protein expression and affinity purification

A maltose binding protein-CahR fusion (MBP-CahR) was engineered in the pMALc2x vector as previously described ^(13), (25), (26)^. Briefly, the coding region of *cahR* was amplified with primers PP_3665Exp2ndMet_EcoRI (5’- AAAGAATTCGTCCACCCCGCTTCTGCAAAC-3’) and PP_3665ExpR_HindIII (5’- AAACAAGCTTAATGCCAATACCTGACCCAA-3’). The ~1 kb fragment was then digested with restriction enzymes EcoRI and HindIII (New England Biolabs, Ipswitch MA), purified using Fisher’s GeneJET Gel Extraction and DNA Clean Up Kit (ThermoFisher Scientific, Waltham, MA), and ligated into the similarly cut pMALc2x vector downstream of the MBP coding region. Cloning and propagation of the pMALc2x:MBP-CahR vector was carried out in *E. coli* DH5α cells grown in LB broth supplemented with 150 μg/ml carbenicillin. The pMALc2x:MBP-CahR vector was then transformed into *E. coli* T7 cells, a strain engineered to support protein expression.

Expression and purification of MBP-CahR was conducted as previously described for MBP-tagged GATR family regulators ^(13), (25), (26)^. Briefly, *E. coli* T7 cells carrying the pMALc2x:MBP-CahR plasmid were grown in a 50 ml volume of LB broth supplemented with 150 μg/ml carbenicillin, shaking at 170 rpm at 37 °C for 3 hours. Protein expression was induced by addition of isopropyl-β-D- thiogalactopyranose (IPTG) to 1 mM followed by an additional 3 hours of growth. Cells were collected by centrifugation, frozen at −20 °C overnight, thawed, and brought up in 4 ml lysis buffer per 1 gram of cells (lysis buffer: 20 mM Tris HCl pH 7.4, 1 mM EDTA, 200 mM NaCl, 1x Halt protease inhibitor (Thermo), 3 mg/ml lysozyme). After a 20-minute incubation to allow for lysis, an additional 5 volumes of lysis buffer was added to further dilute the lysate. Cell lysates were clarified by centrifugation (20 min, 21,000 x g at 4 °C) and applied to an amylose resin column. After application of lysate, the column was washed with 10x the resin bed volume with wash buffer (20 mM Tris-HCl pH 7.4, 200 mM NaCl, 1 mM EDTA) and the protein was eluted using one resin bed volume of elution buffer (20 mM Tris-HCl pH 7.4, 350 mM NaCl, 1 mM EDTA, 10 mM maltose). Expression and purification of MBP-CahR fusion protein was visualized via Coomassie staining of SDS-PAGE gels and protein concentration of the eluate was measured by UV absorbance on a Nanodrop spectrophotometer. Aliquots of protein were stored at −80 °C in 20% glycerol for future use in electrophoretic mobility shift assays (EMSAs).

### Electrophoretic Mobility Shift Assays (EMSAs)

Electrophoretic mobility shift assays (EMSAs) using MBP-CahR protein and a biotinylated promoter probes were conducted as previously described for related GATR family regulators ^(25)^. The promoter regions of several genes predicted to be involved in creatine metabolism were amplified with 5’-biotinylation using the primers noted: *creA/PP_3665* (CreatinaseProm_F_biotin 5’-5’/Biosg/GGTCTTGGGCATTTGCATGG-3’ and CreatinaseProm_R 5’-GTTTGTCCGAGACTTTGTGGC-3’); *glyA1/PP_0297* (PP_0297_F_biotin 5’-5’/Biosg/CAGTACGGAACGGGTCGTAT-3’ and PP_0297_R 5’- GGGTAGCTGCTAGGCTCAAA-3’); and *tdcG-I/PP_0322* (PP_3022_F_biotin 5’- 5’/Biosg/AAACCATCGATTCAGCACTTG-3’ and PP_3022_R 5’- CCTTTGTGGCGATGTTATGA-3’). An additional probe consisting of the promoter region of the *atoA/PP_3122* gene, which is unrelated to creatine metabolism, was tested as a negative control of CahR-promoter binding and was amplified with 5’-biotinylation using primers PP_3122_F_biotin (5’-5’/Biosg/CTGGGCGAAGCTCTGGTACT-3’) and PP_3122_R (5’-TGTTTAACCGACGAGGCTGT-3’). All probes were gel purified using GeneJET Gel Extraction and DNA Clean Up Kit (ThermoFisher). EMSA binding reactions were conducted with the changes previously described with 1 fmol/μl of probe and the replacement of poly(dI-dC) with salmon sperm DNA to a final concentration of 500 μg/ml ^(25)^. Binding reactions were incubated for 40 minutes at 37 °C then run on a precast 5% TBE acrylamide gel at 4 °C for 45 minutes at 100 V (Bio-Rad). After transfer to Biodyne B Modified Nylon Membrane (ThermoFisher), biotinylated probe was visualized as per manufacturer’s instructions with changes as previously described using the Pierce Lightshift Chemiluminescent EMSA kit (ThermoFisher Scientific) ^(25)^.

### Bacterial growth conditions and RNA preparation for RNA-Seq

*P. putida* KT2440 wild type and Δ*cahR* overnight cultures were grown in 1x MOPS amended with 25 mM pyruvate and 5 mM arginine at 30 °C with shaking at 170 rpm. Overnight cultures were washed in 1xMOPS and adjusted to OD_600_ = 1.0 in 1xMOPS with 20 mM pyruvate, and 600 μl of adjusted culture was added to 600 μl pre-warmed 1xMOPS with 20 mM pyruvate +/− 2 mM creatine in a 24-well plate. Each *P. putida* strain was incubated in the +/− creatine condition in technical duplicate and biological triplicate. Cultures were incubated at 30 °C with shaking with sample collection at 1 hour via centrifugation and resuspension in 600 μl of ~60 °C RNAzol RT (Sigma-Aldrich). After lysing cells by pipetting and vortexing in RNAzol, samples were stored until processing at −80 °C. RNA was extracted and purified from these frozen samples using the RNeasy Mini Extraction Kit (Qiagen) as per the manufacturer’s instructions with the following adjustments: after the initial RNA extraction, the RNA samples underwent a DNaseI treatment and an additional sequential RNeasy purification.

### RNA-Seq library preparation

Purified total RNA samples were depleted of rRNA using the MICROBExpress Bacterial mRNA Enrichment Kit protocol (Thermofisher), concentrated via precipitation and resuspension, and mRNA concentrations measured via BioAnalyzer. Precipitated and depleted mRNA samples were used for construction of Illumina-compatible single-end libraries using the NEXTflex Rapid Directional mRNA-Seq Bundle - Barcodes 1-24 (BIOO Scientific). Barcoded libraries were submitted to the Vermont Genetics Network (VGN) sequencing facility at the University of Vermont for generation of read counts via the Illumina HiSeq sequencing system. An average of 11.3 million reads per sample were generated on a HiSeq 1500/2500 single-end 85 bp run.

### RNA-Seq data processing and analysis

Quality assessment of raw sequencing data was performed using FastQC (v0.11.6). Adapters and low-quality sequences were removed using Trim Galore! (v0.6.4) removing Illumina adapters, reads <Q20, and a minimum length of 35 bp. Transcript quantification was performed using Rockhopper2 using the default parameters with verbose output. Reads from each sample were mapped to the reference genome of *Pseudomonas putida* strain KT2440 pre-packed with the program.

Differential abundance was calculated using DESeq2 by Group, a feature encompassing genotype and treatment (i.e., WT Creatine treatment vs WT without Nitrogen). Raw counts were adjusted for library size and genes with fewer than 10 counts in at least 2 samples were removed from further analysis. Normalization was performed using the default settings of DESeq2 with independent filtering and alpha (FDR) set to <0.05. Genes displaying greater than a 2-fold log2 change in transcript levels between conditions were considered differentially expressed and those with a *p*-value of less than 0.05 were considered significant. Gene expression data is available in the NCBI GEO database under accession GSE163362.

### Phylogenetic Tree Building

The amino acid FASTA sequences of orthologs of CreA (creatinase) and associated CahR orthologs (GATR-subfamily AraC members) were compiled via the STRING protein database and BLAST protein searches conducted using the National Centers for Biotechnology Information database ^(27), (28), (29), (30), (31), (32), (33), (34), (35), (36), (37, 38)^. Sequences were entered into a phylogenetic tree-building pipeline available on phylogeny.lirmm.fr (39). This pipeline uses FASTA protein sequences to create a neighbor joining phylogenetic tree using MUSCLE for sequence alignment and PhyML software for tree building. The sequences of a creatinase and GATR from *Pyschrobacter* sp. 4Dc were used as the out-groups for their respective trees due to their distance in similarity from the majority of sequences analyzed. The accession numbers for the amino acid FASTA sequences used for CreA ortholog tree building are: WP_193834191.1, WP_194270409.1, WP_086918772.1, WP_035989123.1, WP_061177555.1, WP_006415961.1, WP_060239820.1, WP_151048639.1, WP_130136211.1, EXE17292.1, WP_187119852.1, WP_077520507.1, WP_090349497.1, WP_043248253.1, WP_147810201.1, WP_005244916.1, WP_005244916.1, ENW47798.1, EXC04443.1, KCY15186.1, KCY60723.1, ENV39029.1, AJB50085.1, EXE17292.1, EXE77507.1, EKA71974.1, EXB17260.1, EXC10249.1, WP_001094923.1, WP_001094923.1, AAD52565.4, AEJ12833.1, AHC82230.1, AHC87608.1, AHZ77043.1. Accession numbers for the protein FASTA sequences of CahR orthologs associated with the above CreA sequences are: WP_101206352.1, WP_153168678.1, WP_051495264.1, WP_051453741.1, WP_061177577.1, WP_048994731.1, WP_060239822.1, WP_151048638.1, WP_004832906.1, EXE17291.1, WP_058356168.1, WP_077520504.1, WP_090349496.1, WP_043248255.1, WP_147810202.1, WP_005244913.1, WP_005244913.1, ENW47797.1, EXC04444.1, KCY15187.1, KCY60724.1, ENV39028.1, AJB50136.1, EXE17291.1, EXE77506.1, EKA71975.1, EXB17259.1, EXC10250.1, WP_000941154.1, WP_000941154.1, AEJ12829.1, AHC82227.1, AHC87605.1, AHZ77039.1.

## RESULTS

### Identification of *PP_3665* (*cahR*) as essential for *P. putida* KT2440 utilization of creatine as a sole nitrogen source

The location of the uncharacterized GATR *PP_3665* near the *creA* gene led to the prediction that it would function to regulate creatine metabolism in *P. putida*. To test this prediction, we evaluated wild type and the Δ*PP_3665* deletion strain’s abilities to grow on various nitrogen sources related to creatine metabolism. After 18 hours of incubation *P. putida* wild type, *P. putida* Δ*PP_3665*, and the complemented strain all grew equally efficiently on choline, arginine, and sarcosine as sole nitrogen sources (**Fig. 1C**). Growth of *P. putida* Δ*PP_3665* on creatine was significantly lower when compared to WT and complemented strains (p <0.0001), showing no net growth compared to the no-nitrogen control media (**Fig. 1C**). When supplied with creatinine, the anhydrous form of creatine, as the sole source of nitrogen, all strains had lower growth compared to their growth in choline or to wild type in creatine (p < 0.01), but growth of the *P. putida* Δ*PP_3665* strain was not different than wild type in creatinine (**Fig. 1C**), suggesting general poor growth is likely due to inefficient creatinine utilization and potentially an alternate route in this strain of *P. putida*. Based on its essential role in creatine-dependent growth and as a transcription regulator, we named *PP_3665* as *cahR* (**<u>c</u>**reatine **<u>a</u>**mino**<u>h</u>**ydrolase **<u>r</u>**egulator) and use that nomenclature for the remainder of this report.

### CahR induces *creA* transcription in the presence of creatine

Compounds related to the creatine metabolic pathway were tested for their ability to induce *gfp* in a *cahR*-dependent manner from a *creA* promoter-*gfp* fusion. Significant transcriptional induction was observed in the presence of creatine, ~13 fold over the no-inducer condition (p <0.0001) (**Fig. 2A**). Induction of the *creA* promoter in the presence of creatine was also significantly higher in wild type compared to the Δ*cahR* deletion strain, in which fluorescence was similar to the no-inducer condition, indicating that creatine-dependent transcription induction from the *creA* promoter is *cahR*-dependent and CahR functions as a transcriptional activator.

**Figure 2.**
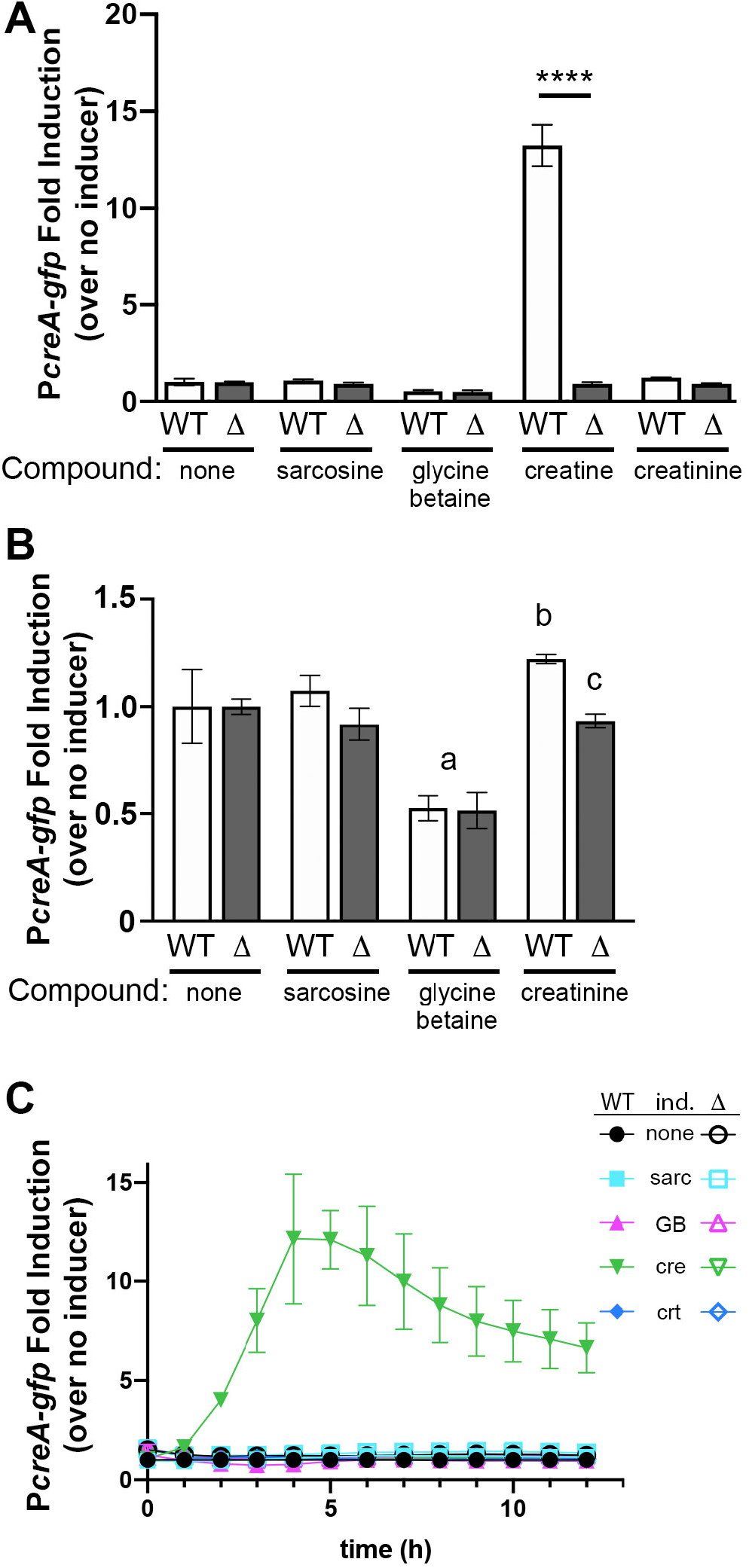
Induction of a GFP transcriptional reporter to the *creA* promoter region. **(A)** Creatine strongly induces the *P_creA_-gfp* reporter in wild-type cells (WT) but not in the Δ*cahR* strain (Δ). Statistical significance tested using ANOVA with Sidak’s post-test comparing wild-type to deletion within each condition. The four asterisks represents p < 0.0001; all other statistical comparisons are note in panel B. **(B)** Data replotted from (A) but leaving out the creatine condition to emphasize small but replicable changes driven by glycine betaine and creatinine. Statistical analysis tested using ANOVA with Sidak’s post-test comparing all pairs of data. The glycine betaine condition represses expression significantly independent of *cahR* (a, denotes p < 0.001 in comparison to no inducing compound). Creatinine very slightly but replicably induces the reporter (b, denotes p < 0.05 in comparison to no inducing compound), which is dependent on *cahR* (c, denotes p < 0.05 in comparison to WT creatinine). **(C)** Timecourse of induction from the *creA* promoter in wild type (WT, filled symbols) and Δ*cahR* (Δ, open symbols) in the presence of no inducer (none), sarcosine (sarc), glycine betaine (GB), creatine (cre), and creatinine (crt). Error bars in all panels represent standard deviation.

A closer look at induction of the *creA* promoter in the weakly-inducing conditions shows that glycine betaine represses expression independent of *cahR*, while creatinine mildly induces the reporter in a *cahR*-dependent manner (**Fig. 2B**). The creatinine data suggest that creatinine might interact with CahR poorly but in a manner that stimulates transcriptional induction or that our commercial creatinine has trace amounts of creatine contamination. Glycine betaine suppression of the *creA* promoter is similar to GbdR-dependent suppression of alternate GATR-controlled loci in *P. aeruginosa* ^(13, 40)^.

The transcriptional response of the *creA* promoter to creatine is specific and rapid as assessed during a fluorescence time course. *creA* reporter induction is only seen in WT in the presence of creatine, with normalized reporter activity peaking about four hours after addition of creatine (**Fig. 2C**). It is important to note that the activity of the reporter lags native *creA* transcript accumulation, which we assessed by RT-PCR as peaking roughly at one hour post induction (data not shown).

### CahR binds the upstream regulatory region of *creA*

MBP-CahR binds to the promoter region of *creA*, shifting the *creA* promoter probe in a concentration-dependent manner, but not substantially shifting the non-specific *P. aeruginosa atoA* promoter probe or the creatine metabolism-related genes *glyA-1* and *tdcG-I* (**Fig. 3**). GATR family regulators are often poorly soluble and while we have purified and examined some without epitope tagging ^(40, 41)^, fusion of maltose-binding protein (MBP) to the amino terminus greatly enhances solubility and does not alter DNA binding site specificity for other GATR family members ^(13, 25, 40–42)^. Additionally, from those same studies and including those without epitope tags, ligand binding does not alter GATR association with DNA, a property we also confirmed with CahR (data not shown).

**Figure 3.**
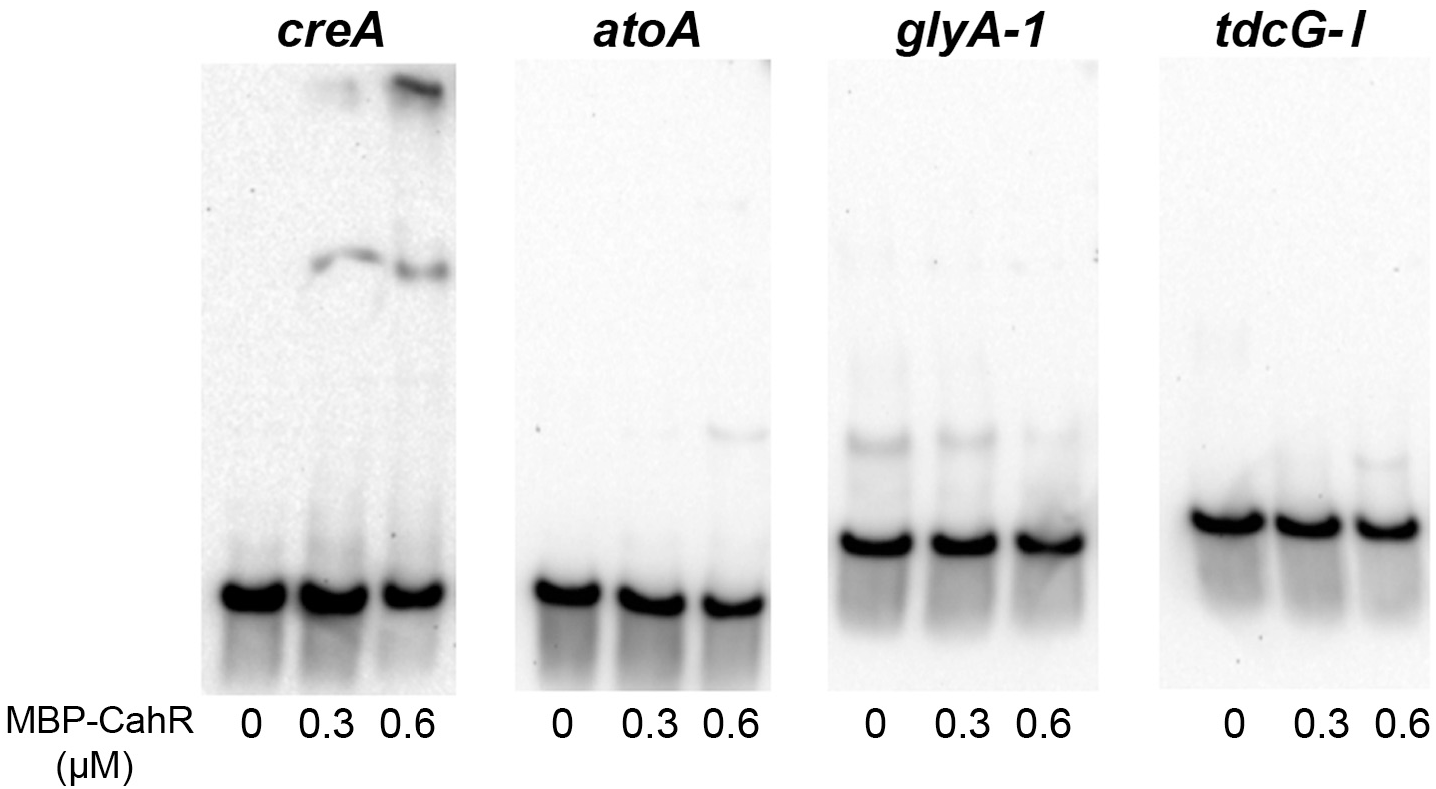
Electrophoretic mobility shift assays (EMSA) with purified MBP-CahR. Purified MBP-CahR (concentrations noted at bottom of each lane) was incubated with the biotinylated promoter probes labeled at the top of each blot. Strong and specific shift only noted with the *creA* promoter.

### Identification of the CahR binding site in the *creA* promoter

*cahR*-dependent transcriptional induction of the *creA* promoter is highest when the full promoter region (between −10 bp and −195 bp from the predicted transcriptional start site) is present. When the promoter is truncated to include only −169 bp upstream of the predicted transcriptional start, transcriptional induction drops substantially compared to the full-length construct and is only ~40% higher than the uninduced condition (**Fig. 4A**). Truncations of the *creA* promoter to −80 bp or beyond eliminate all creatine-dependent transcriptional induction of the *creA* reporter. The creatine-dependent induction of the −195 bp and −169 bp reporters is also *cahR*-dependent, with a significant difference in fold induction observed between the reporters in the *P. putida* wild type and Δ*cahR* strains (−195 bp p < 0.0001, −169 bp p < 0.001) (**Fig. 4B**).

**Figure 4.**
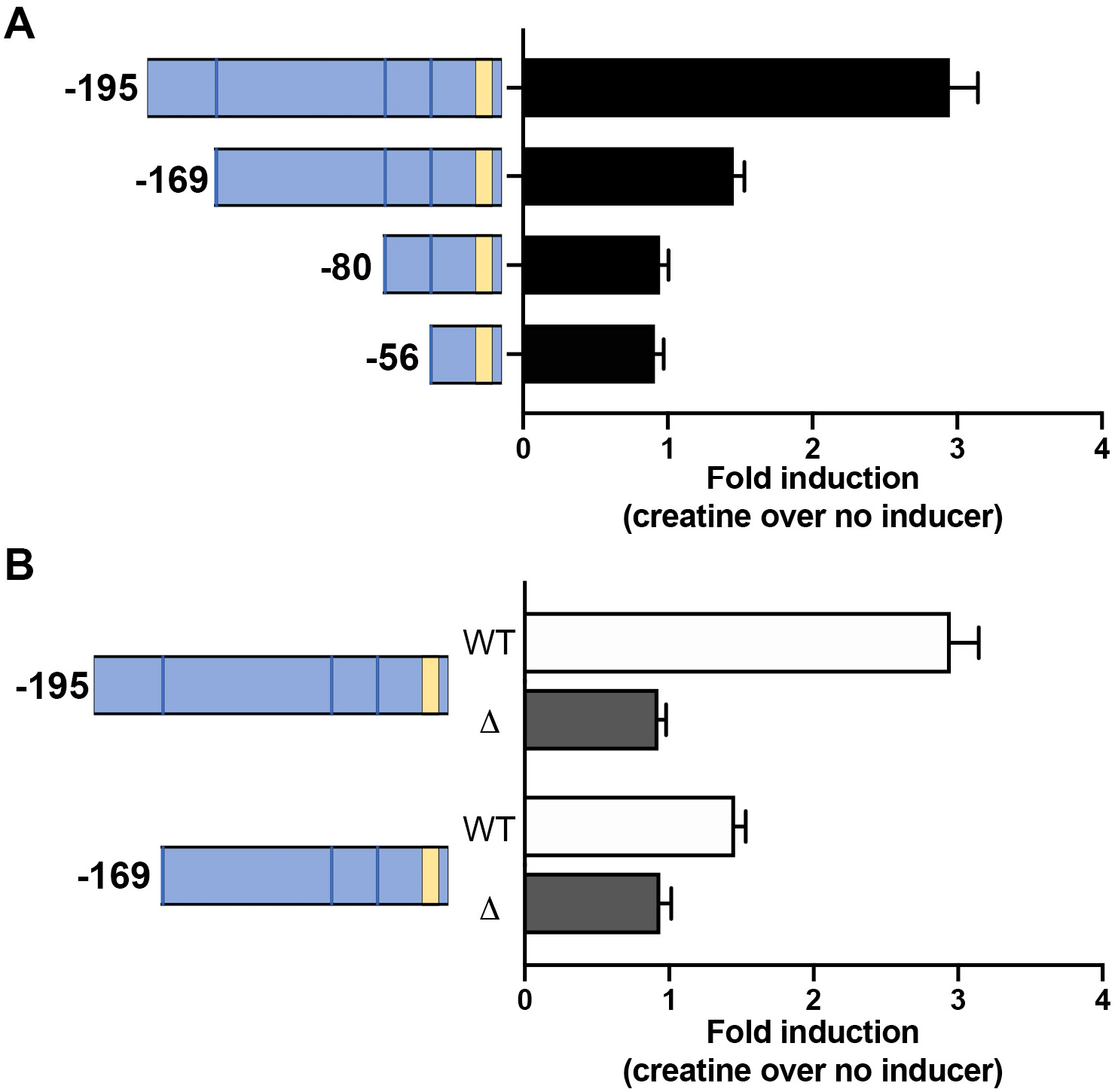
Mapping the likely CahR binding site in the *creA* promoter. Promoter truncations based on the *P_creA_-gfp* reporter were used to assess the minimal fragments that retain creatine/*cahR*-dependent induction. **(A)** Fold induction of four promoter truncations in wild type cells. **(B)** Expression from the two largest promoter truncations in wild-type (WT) and the *cahR* deletion (□). In both panels, tan block represents the predicted promoter.

### CahR is required for transcription of creatine and sarcosine metabolism-related genes in *P. putida* in response to creatine

To determine the genes involved creatine utilization by *P. putida*, wild type was exposed to 2 mM creatine or 0 mM creatine in nitrogen-free minimal media for 1 hour. There were 22 transcripts differentially induced more than 4-fold in the 2 mM creatine condition compared to the 0 mM creatine control condition, the majority of which are predicted to be involved in creatine and sarcosine metabolism (**Fig. 5 and Table 1**). The gene with the highest induction over the control condition, 1260-fold, is *PP_3667/creA* which encodes the known *P. putida* creatinase. The other member of the *creA*-containing operon, *PP_3666* encoding a putative metabolite MFS transporter, was also among the creatine-responsive genes, induced 362 fold (**Fig. 5A and Table 1**).

**Figure 5.**
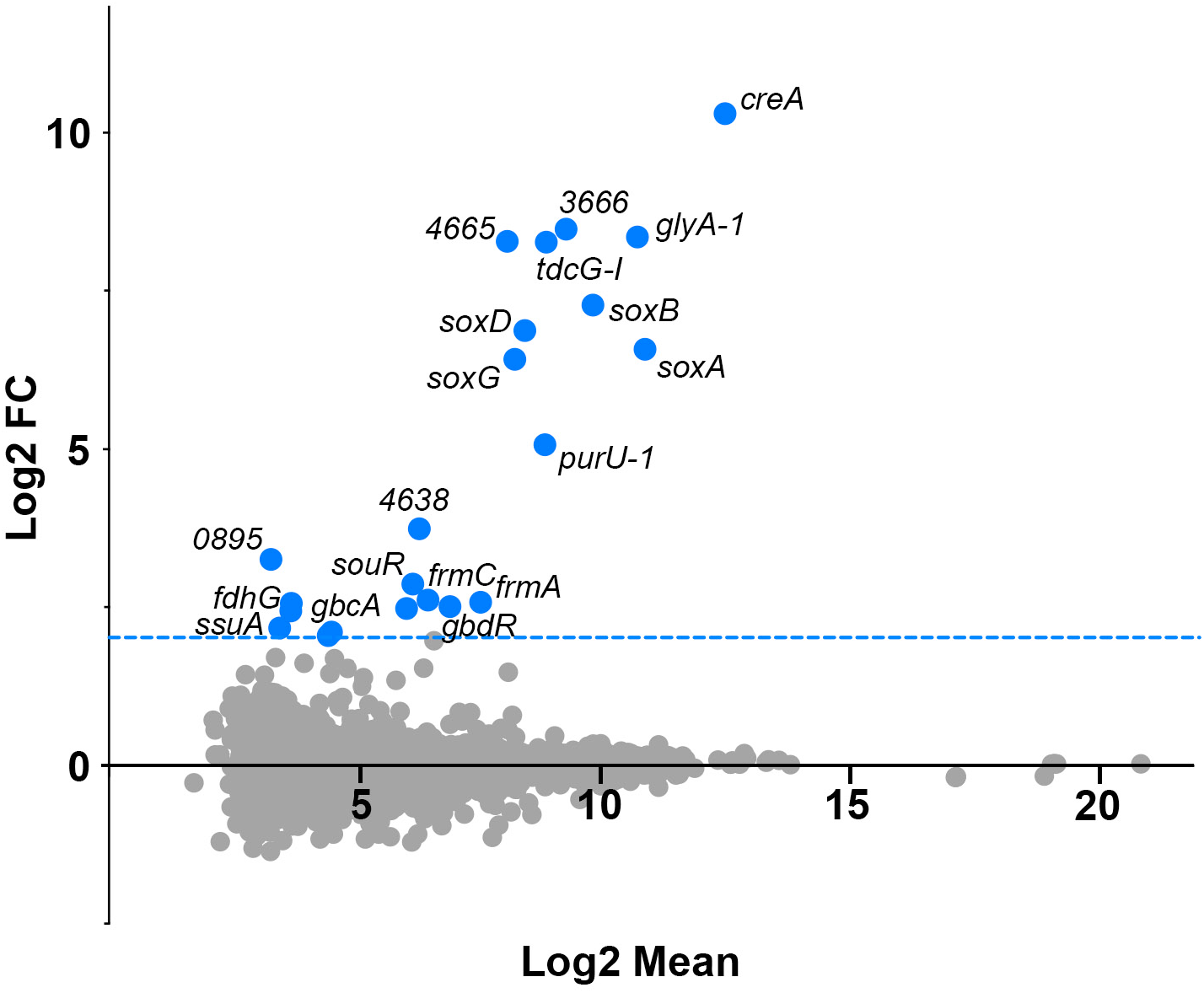
Mean-difference (MD) plot of transcript abundance comparing wild type in creatine over wild type with no added nitrogen source. Wild type *P. putida* KT2440 was transferred to media with or without creatine as a sole nitrogen source for 1h, after which RNA was harvested and transcriptomics by RNA-Seq conducted. Transcripts that met the fold change (Log2 > 2) and p-value (p < 0.05) are shown in blue and labeled where space allowed. All transcripts meeting these two qualifications are listed in **Table 1**.

**Table 1.** Differentially expressed transcripts in wild-type cells in the absence and presence of creatine.

| Gene | log2FoldChange | log2 Mean | Product Function / Product Relationship |
| --- | --- | --- | --- |
| PP_3667 *creA* | 10.3 | 12.5 | creatinase |
| PP_3666 | 8.5 | 9.3 | major facilitator superfamily transporter |
| PP_0322 *glyA-1* | 8.3 | 10.7 | serine hydroxymethyltransferase |
| PP_4665 | 8.3 | 8.1 | PA2762 ortholog |
| PP_0297 *tdcG-1* | 8.3 | 8.9 | L-serine dehydratase |
| PP_0323 *soxB* | 7.3 | 9.8 | sarcosine oxidase subunit beta |
| PP_0324 *soxD* | 6.9 | 8.4 | sarcosine oxidase subunit delta |
| PP_0325 *soxA* | 6.6 | 10.9 | sarcosine oxidase subunit alpha |
| PP_0326 *soxG* | 6.4 | 8.2 | sarcosine oxidase subunit gamma |
| PP_0327 *purU-1* | 5.1 | 8.8 | formyltetrahydrofolate deformylase |
| PP_4638 | 3.7 | 6.3 | methylenetetrahydrofolate reductase domain-containing protein |
| PP_0895 | 3.3 | 3.3 | hypothetical protein |
| PP_3526 | 2.9 | 6.2 | Predicted SouR ortholog |
| PP_1617 *frmC* | 2.6 | 6.5 | S-formylglutathione hydrolase |
| PP_1616 *frmA* | 2.6 | 7.5 | D-isomer specific 2-hydroxyacid dehydrogenase |
| PP_0896 | 2.6 | 3.7 | C/N hydrolase; nitrilase/cyanide hydratase |
| PP_0299 | 2.5 | 6.9 | Predicted GbdR ortholog |
| PP_3543 | 2.5 | 6.0 | (Fe-S)-binding protein |
| PP_2183 *fdhG* | 2.4 | 3.7 | formate dehydrogenase subunit gamma |
| PP_0236 *ssuA* | 2.2 | 3.5 | NAD(P)H-dependent FMN reductase |
| PP_0315 *gbcA* | 2.1 | 4.5 | Rieske (2Fe-2S) domain-containing protein; GB metabolism |
| PP_0894 | 2.1 | 4.4 | NTF2 family protein |

It was not surprising that the creatinase gene was the most highly expressed gene in the presence of creatine, as lysis of creatine into urea and sarcosine is generally the first step in bacterial creatine metabolism. The predicted pathway of *P. putida* creatine metabolism is outlined in **Fig. 1A** and is supported by our differential expression data. Following the hydrolysis of urea from creatine, the resulting sarcosine molecule is oxidatively demethylated into glycine and formaldehyde by the tetrameric sarcosine oxidase encoded by *soxBDAG*. The *sox* operon *soxBDAG* genes are differentially induced between 64 and 128 fold in 2 mM creatine over the control condition. The glycine that results from sarcosine oxidation can then be converted to serine via the glycine hydroxymethyltransferase encoded by *glyA-1*, which is expressed approximately 256 fold in 2 mM creatine over the control condition. Finally, serine can be converted into pyruvate via serine dehydratase encoded by the *tdcG-I* gene (ortholog of *P. aeruginosa sdaB*) that is expressed 256 fold higher in the 2 mM creatine as compared to the control. The pyruvate generated from creatine metabolism is then available for conversion into acetyl-CoA and thus into central metabolism. Taken together, the induction of creatine and sarcosine-metabolic genes provides support for the previously predicted pathway of creatine metabolism in *P. putida* KT2440 (**Fig. 1A)**.

The role of CahR in creatine-responsive gene induction was also elucidated using RNA-seq and differential expression analysis. The 2 mM creatine versus 0 mM creatine comparison was repeated as above, but with the Δ*cahR* mutant. When the Δ*cahR* mutant was exposed to 2 mM creatine, the creatine and sarcosine-metabolic genes were no longer differentially expressed over the control condition, leaving only a single gene that met the cut-off criteria used for wild type – the nitrogen fixation-related gene, *fixG*. The lack of creatine metabolic gene induction in the absence of *cahR* indicates that CahR is responsible for the transcription of these genes. The lack of induction of the genes encoding downstream metabolic steps, including sarcosine oxidation in the absence of evidence for their direct control by CahR (see **Fig. 3**), is not surprising as production of sarcosine is dependent upon a CahR-regulated step. We also confirmed that creatine does not induce a *sox* operon transcriptional reporter in the absence of *creA* (data not shown). Complete RNA-Seq data is available at (NCBI GEO Accession currently in submission).

### *creA* orthologs associated with a *cahR* ortholog are scattered throughout the □- and □-proteobacteria

Orthologs of the *creA* creatinase cluster into two clades, one with the *P. putida* and related Pseudomonads and the other with ß-proteobacteria and *Acinetobacter* (**Fig. 6**). A number of *Acinetobacter* species maintain a genomic region that is orthologous to *P. knackmussii*, *P. oryzae*, and Burkholderia creatinase-coding regions but is present flanked by transposable element boundaries. For some strains this creatinase-containing transposon is on the chromosome whereas in three others, it is plasmid-borne. The presence of creatine-metabolic genes on a transposon maintained in pathogenic bacteria suggests that creatine metabolism may be beneficial during pathogenesis. The presence of creatinase and related metabolic genes in both environmental and pathogenic species suggests a role in both niches, but the strain specific carriage of these genes and alternate gene organization in otherwise closely related species strongly suggests that these genes are readily acquired via horizontal gene transfer.

**Figure 6.**
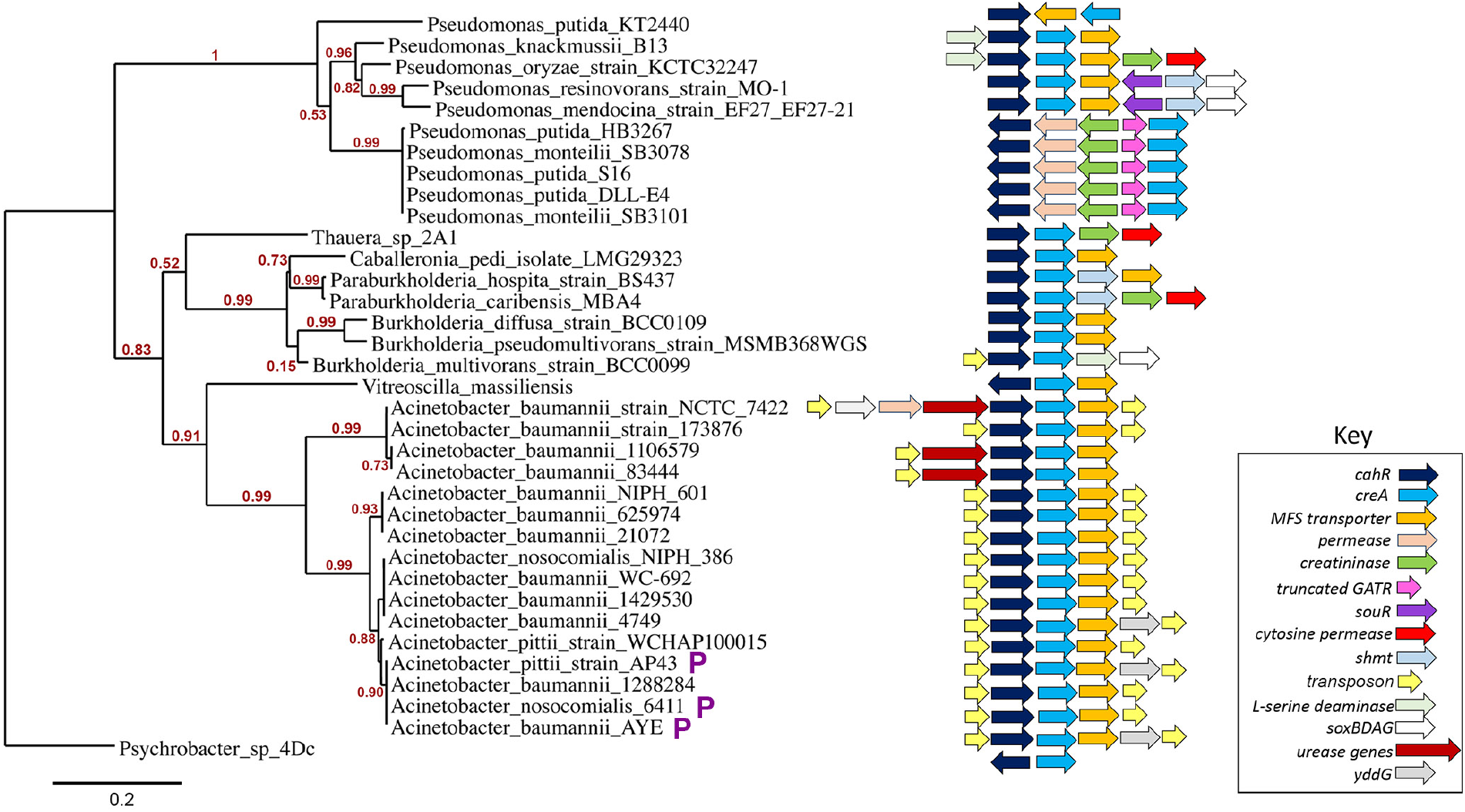
Phylogenetic tree of CreA amino acid sequence with associated genomic context of the *creA* gene. Predicted amino acid sequences from *creA* genes were examined for nearby *cahR* orthologs and resultant CreA protein sequences were phylogenetically analyzed using MUSCLE alignment of amino acid sequences and PhyML tree construction. Numbers shown next to branches are approximate likelihood ratio scores. Predicted function for genes in the *creA* genomic regions are denoted by color as shown in the key. Abbreviations: serine hydroxymethyl transferase, *shmt*; transposable element components including transposases and transposable element flanking sequences, *transposon*; genes present on plasmids in this strain, P in purple text.

## DISCUSSION

Creatine is a nitrogen-rich compound (N:C ratio 1:1) present in many of the ecological niches occupied by *Pseudomonas* species. The best described creatine pool is found within vertebrate muscle cells, where it participates in the rapid recharging of ADP to ATP through a direct phosphate transfer cycle. The body is constantly recycling its creatine stores and excreting creatine into the surrounding environment through urine and feces ^(15), (16), (17), (18)^. Most of the creatine in the body is non-enzymatically converted into creatinine before excretion, making this cyclic form of creatine abundantly present in the colon, where gut-resident microbes degrade it, and the rhizosphere where it can be utilized by soil-dwelling microorganisms ^(18), (43), (44), (45)^. Because of the conversion of creatine to creatinine within vertebrates before excretion, many bacterial pathways for creatine metabolism begin with lysis of creatinine via creatinine amidohydrolase enzymes ^(45), (46), (47), (48)^ (**Fig. 1A**). However, the existence of creatine-responsive metabolic genes in primarily soil dwelling bacteria that lack a creatininase suggest that creatine is available within the rhizosphere for utilization. Free creatine may come from a variety of sources, including creatinine breakdown by creatininase-possessing microbes, the in situ degradation of creatine-containing animal tissues, or excretion of smaller amounts of creatine in animal urine. Regardless of the source, the ability to metabolize creatine would enable bacteria to access a rich nitrogen source and in some cases an alternative source of carbon ^(10), (49)^. This manuscript describes identification of a creatine-responsive transcription regulator, CahR, in *P. putida* that is critical for utilization of creatine as a sole nitrogen source.

### *cahR* is essential for creatine-dependent induction of *P. putida* creatine utilization genes, thus growth on creatine as a sole nitrogen source

The AraC/XylS-family of transcriptional regulators are a diverse family generally characterized by a helix-turn-helix (HTH) DNA binding C-terminal domain and an N-terminal domain dedicated to dimerization and/or ligand binding ^(50)^. The creatine-responsive transcriptional regulator of *P. putida*, CahR, belongs to a subset of AraC/XylS-family regulators that contain a glutamine amidotransferase-1 (GATase) domain in the N-terminal region with structural similarity to the DJ-1/ThiJ/PfpI superfamily of proteins. Members of this superfamily include the human DJ-1 protein, involved in muscular dystrophy^(51, 52)^, and the bacterial ThiJ proteins involved in bacterial thiamine biosynthesis ^(53)^. Bacterial AraC-family transcriptional regulators in the GATase subfamily have a predicted catalytically-inactive ThiJ-like domain in the N-terminus, but that often includes a conserved cysteine residue found in the catalytic domain of active bacterial GATase proteins such as *Klebsiella pneumoniae*’s ThiJ ^(54), (41), (53)^. Members of the GATase 1-containing AraC transcriptional regulator (GATR) family have been identified in multiple Gram-negative and Gram-positive bacteria, including *Pseudomonas* species, although functions have only been described for a limited subset of these regulators. Several of the characterized GATRs in *P. aeruginosa* participate in metabolic regulation of amine-containing compounds like arginine, glycine betaine, sarcosine, and carnitine ^(13), (25), (55), (56), (57, 58)^. The *P. putida* GATR described in this manuscript, CahR, controls the metabolism of the amine-containing compound creatine via transcriptional induction of the creatinase CreA.

The participation and necessity of *cahR* in creatine metabolism is demonstrated in **Fig. 1C**, where deletion of *cahR* results in the inability of *P. putida* to grow on creatine as the sole nitrogen source. CahR binds with specificity to the promoter region of *creA* (**Fig. 3**), from which we conclude direct transcriptional induction of *creA*, which encodes a creatinase with the ability to efficiently cleave creatine into sarcosine and urea ^(5), (6), (7), (8), (9), (59), (60), (61)^. Transcription of *creA* occurs quickly in wild type *P. putida*, detectable by GFP reporter within 1 hour after exposure to creatine, and with rapid increase over the first 5 hours post-exposure (**Fig. 2C**). At one hour post creatine exposure, a ≈1260 fold-change in transcript levels of *creA* is observed in *P. putida* WT in the presence of creatine versus a pyruvate control (**Fig 5** and **Table 1**). This suggests that lysis of creatine by CreA is the preferential pathway of *P. putida* creatine metabolism and the rapidity of creatine metabolic induction compared to rates for other GATRs suggests that creatine utilization is likely a beneficial metabolic strategy for *P. putida* and/or that creatine is a resource under strong competition.

The metabolism of creatine by *P. putida* creatinase to the intermediate sarcosine has been observed by multiple groups and is supported by the strong transcriptional induction of the predicted sarcosine metabolic genes, orthologous to *Pseudomonas aeruginosa’*s *soxBDAG*, one hour post creatine exposure in *P. putida* KT2440 (**Fig. 5** and **Table 1**) ^(5), (6), (7), (8), (9), (59), (60), (61)^. Genes involved in the subsequent steps of sarcosine metabolism, including *glyA1* encoding the serine hydroxymethyltransferase and *tdcG-I* encoding the L-serine dehydratase, are amongst the next most highly transcribed genes in the presence of creatine (**Fig 5** and **Table 1**), providing a more complete picture of creatine metabolism in *P. putida* KT2440, as outlined in **Fig. 1A**. Although multiple genes are induced in the presence of creatine, CahR specifically binds to the promoter region of *creA* alone and not to the promoter regions of the other metabolic genes most highly induced in the presence of creatine, suggesting a small creatine-specific regulon controlled by CahR (**Fig. 3**). Based on promoter mapping, CahR’s specific binding site lies within region −195 bp to −169 bp from the predicted transcriptional start of *creA* and likely close to or partially overlapping the −169 position. Unfortunately, CahR’s apparent specificity for a single promoter prevented further prediction of a specific CahR binding site, as there is no additional promoter(s) bound by CahR to use in identifying conserved half site sequences. We did attempt alignments and motif detection between strains and species and also did not identify a potential conserved CahR binding site.

While *P. putida* KT2440 is able to utilize creatine and the downstream metabolite sarcosine as sole nitrogen sources, it grows poorly on creatinine (**Fig. 1C).** This is interesting, as creatinine is generally considered a precursor to creatine in the context of bacterial metabolism (**Fig. 1A**). However, creatine may or may not be an intermediate in creatinine metabolism in *P. putida* KT2440, as there are alternatives in some *P. putida* strains including creatinine metabolism via N-methylhydantoin and N-carbamoylsarcosine intermediates^(46)^. Thus, the poor *P. putida* KT2440 growth on creatinine as a sole nitrogen source, independent of CahR-dependent creatinase induction, may be due to inefficient creatinine utilization via an intermediate that is not creatine ^(46), (62)^. The hypothesis that creatinine, when available, is converted by *P. putida* KT2440 into an intermediate that is not creatine, such as N-methylhydantion, is also supported by the negligible induction of *creA* in the presence of creatinine (**Fig. 2A-B**). Creatinine metabolism by a non-creatine intermediate is also supported by the observation that *P. putida* KT2440 does not appear to encode any predicted creatinine amidohydrolases, while several other strains of *P. putida*, including strain S16 (CP002870.1), DLL-E4 (CP007620.1), HB3267 (CP003738.1), and RS56 (AF170566.3) encode creatininases transcribed divergently from *cahR* and *creA* orthologs (**Fig. 6**). The evidence for creatininase function was demonstrated using the cloned and purified enzyme from *P. putida* strain RS56 ^(63)^. In addition to the *P. putida* strains that encode both creatininases and creatinases, there are several *P. monteilii* strains, a close relative of *P. putida*, which share syntentic creatininase/creatinase genomic regions, including *P. monteilii* SB3101 (CP006979.1) and SB3078 (CP006978.1). *P. monteilii* has been implicated in several opportunistic infections, while *P. putida* HB3267, a strain isolated from hospitalized patients and also contains this gene arrangement, shows cytolytic activity against human cells^(64),(65)^.

It is also interesting to note that glycine betaine inhibits basal transcription of *creA* independent of CahR. This may be indicative of an inhibitory feedback mechanism perpetuated by downstream products of creatine metabolism or the direct or tangential involvement of other compound-specific regulators in creatine metabolism. In *P. aeruginosa*, the glycine betaine/dimethylglycine sensing regulator GbdR is able to repress activation from promoters co-regulated with other GATR family members when glycine betaine is present, best described at the carnitine operon promoter that is induced in a carnitine-dependent manner by the GATR member CdhR^(40)^. Increased transcription of GbdR and glycine betaine metabolic genes is also observed in wild type *P. putida* KT2440 in the presence of creatine, which supports potential interplay between *P. putida* GATRs (**Fig 5**).

### Conservation of *creA* and *cahR* synteny illustrate the potential utility of this genetic module in creatine rich environments

Pathways for creatinine and creatine metabolism are conserved among diverse bacteria. Examining sequence similarity and genomic organization between bacteria, the presence of a CahR-regulated creatine-inducible creatinase appears to be conserved among several species, based on the presence of *creA* orthologs co-occuring with predicted *cahR* orthologs in similar genomic organization as *P. putida* (genomic organization in *P. putida* KT2440 shown in **Fig. 1B**). The phylogenetic tree in **Fig. 6** is for CreA, while a tree of the associated CahR orthologs has a very similar topology but, as expected for a regulator compared to an enzyme, shows substantially longer branch lengths (data not shown).

The presence of a *creA* ortholog with an associated *cahR* ortholog in a limited number of strains within a given species supports a model of horizontal gene transfer of creatinase-responsive metabolic genes between bacteria. The case of *cahR* and *creA* orthologs among *Acinetobacter* species provides the most compelling support for horizontal transfer of *cahR* and *creA*. The clade with the shortest branch lengths consists exclusively of *Acinetobacter* species, including *A. baumannii, A. nosocomalis*, and *A. pittii*. These *Acinetobacter* species are members of the *Acinetobacter calcoaceticus*-*Acinetobacter baumannii* complex (ACB) and are known for their ability to cause persistent, multidrug resistant, nosocomial infections in humans^(66),(67)^. Of the *Acinetobacter* isolates possessing *cahR*/*creA* orthologs, several were isolated from urine, which is the primary vehicle of creatine excretion in animals^[8]^. Additionally, all of the *Acinetobacter* clinical isolates analyzed possess *cahR*/*creA* orthologs flanked by transposon or integrase flanking sequences, suggesting that the creatine-responsive creatine-metabolic enzymes are part of a transposable element horizontally transferred amongst pathogenic *Acinetobacter* species.

The majority of the *cahR/creA*-containing transposons present in the *Acinetobacter* species are identical, containing *cahR, creA*, and *PP_3666* (encoding the MFS transporter) flanked by transposase/integrase flanking regions, while several species include additional genes related to creatine metabolism such as urease genes, within the putative transposable element (**Fig 6**). In three cases, the *cahR*/*creA*-containing transposon is present on a plasmid that is maintained by *Acinetobacter*. The strains possessing these plasmids, *A. baumannii* AYE (SAMEA3138279), *A. pittii* AP43 (SAMN12612836), and *A. nosocomalis* 6411 (SAMN03263968), cluster together (**Fig 6**). The AP43 plasmid also carries virulence factor *blaNDM-1*, which confers resistance to carbepenems and cephalosporins, while AYE is associated with MDR community-acquired infections^(68)^. The maintenance of the *cahR*/*creA* orthologs, both on plasmids and within transposable elements, suggests that these genes were likely acquired from other sources, such as *cahR/creA*-possessing gut microbiome members *Vitreoscilla massiliensis* or *Thauera* 2A1, or from one of the many environmental or opportunistic pathogen species that share *Acinetobacter’s* niches^(69),(70)^. While the existence of *cahR*/*creA*-containing transposons and plasmids suggest that acquisition and maintenance of these genes is advantageous to *Acinetobacter*’s lifestyle, the potential benefits of this region for bacterial survival and virulence have yet to be evaluated.

### Conclusions

Here we have described the identification of a creatine-responsive transcription regulator, CahR, that is necessary for creatine utilization by regulating creatinase gene induction. There are a number of issues raised by the data presented here that remain to be explored. Based on our count this is now the fourth GATR for which an inducing ligand is known, yet we still do not understand how ligands are bound and how specificity is determined. These GATRs all control organic nitrogen compound utilization and are likely not-so-ancient paralogs that diversified for specialization to structurally related but different small molecules. Thus, the GATRs might provide a good model to understand the evolution of substrate specificity for transcription regulators. Finally, the ecological and/or virulence function for creatine metabolism and its regulation is not understood. The obvious horizontal gene transfer of this metabolic system in Acinetobacter might offer a promising system to test the function of creatine metabolism and regulation in an opportunistic pathogen.

## Acknowledgements

We would like to thank Alexis Nadeau for technical assistance during their research rotation. The next-generation sequencing and bioinformatic analysis was performed in the Vermont Integrative Genomics Resource Massively Parallel Sequencing Facility and was supported by the University of Vermont Cancer Center, Lake Champlain Cancer Research Organization, UVM College of Agriculture and Life Sciences, and the UVM Larner College of Medicine. This work was supported in part by R21AI137453 and internal funding from the Larner College of Medicine to MJW. LAH was supported by T32 AI055402.

